# Can crop phenology and plant height be channelized to improvise wheat productivity in diverse production environments?

**DOI:** 10.1101/2020.10.06.327890

**Authors:** D Mohan, H M Mamrutha, Rinki Khobra, Gyanendra Singh, GP Singh

## Abstract

Non-grain parameters like height, flowering and maturity should also be tried to break yield plateau in wheat. This study explores such possibilities by analysing performance of released and pre-released varieties evaluated in ten diverse production environments of India during the period 2000-2020. Regression analysis supports relevance of such non-grain determinants in grain yield under every environment but magnitude of impact can vary. Collective contribution of non-grain parameters can be high in a production environment where growth condition is most favourable for wheat growth and every factor is important in such situations. They contribute less in the environments engrossed with abiotic stress and merely one or two factors can be earmarked for selection. Besides yield, this selection strategy can also enhance grain weight in certain environments. At a time when selection for grain attributes is not providing further push; it would be worth trying to explore these non-grain field indicators as selection strategy for further advancement in productivity and grain weight of bread wheat.

## Introduction

Wheat (*Triticum aestivum L*.) is grown under diverse agro-ecological conditions across the globe. In India also, it is cultivated under different production environments where growth conditions differ and so is the yield harvest. It is obvious that besides productivity, field expression must also be differing under diverse growth environment (Pandey *et al*., 2015; Sharma *et al*., 2019). Thus, understanding of the changes occurring in the grain and non-grain yield parameters and the inter trait relationship become important for the wheat breeders for further hike in yield potential. Plant phenology which describes the timing of plant development has been acknowledged as a major aspect of plant response to environment. Changing crop phenology can serve as important bio-indicator in the era of climate change (Asseng *et al*., 2017; Rezaei *et al*., 2018). The adapted early flowering cultivars successfully advance the onset of anthesis and the enforced longer grain filling period reduces or avoids the risks of exposure to enhanced drought and heat stresses in late spring (Solanki *et al*., 2017; Yang *et al*., 2019). Optimal height under given environmental condition is vital for adaptability, productivity and yield stability of the wheat cultivars (Bognár *et al*., 2007) whereas maturity duration is the major genotypic cause of genotype-environment interaction (Garatuza-Payan *et al*., 2018; Singh *et al*., 2017; Xie *et al*., 2015).

Traditionally, grain number and grain weight have been recognized as main constituent of wheat yield (Brinton and Uauy, 2019; Garatuza-Payan et al., 2018) and wheat breeding programme emphasise increase in the grain number through better tillering and spike characteristics. In some wheat breeding centres of India, grain weight is also addressed in combination with the heat tolerance programme (Braun et al., 2010; Mishra et al., 2014; Mondal et al., 2016). Still, the yield level keeps staggering and raising of the yield bar even by 5-10% turn out to be a difficult proposition in certain regions. At this juncture, it is imperative to explore the role and contribution of non-grain parameters (NGP’s) namely plant height, maturity duration and heading days. It is a general perception that adversary of climate change in wheat is first realised on NGP’s and later reflected in grain yield and size of the grain. Increased height and crop duration under favourable growth condition often results in higher biomass production and accumulates more grain yield (Reynolds *et al*., 2009). Although NGP’s are influenced by the abiotic factors; genetic constitution also modulates their role in ascribing varietal differences. Therefore, it is crucial to understand whether selection exercised on these field indicators can lead to yield improvement, if so up to what extent and under which environment they can be exploited. Such studies attain more prominence when production environments are highly diverse as observed in India. Few reports from India have highlighted variations in the grain and non-grain attributes of wheat under diverse growth environments (Mohan *et al*., 2011; Mohan *et al*., 2017). However, a comparative study to demonstrate their contribution and potential role in yield enhancement without exerting any undesired effect on grain size was lacking. Yield is expensive to pursue, therefore other objectives must be attained before wide scale yield evaluation. Indian wheat programme provides perfect platform for such investigations where high-yield genotypes of different genetic background are being tested in several productions environments for a long time and data has been generated on plant height, days to heading and maturity duration, grain yield and grain weight. Examining long-term yield data of Indian wheat research programme, this study is an attempt to i) emulate differential impact of NGP’s and understand the interrelationship pattern, ii) realize their comparative contribution in grain yield, iii) suggest ways to tap their potential for further increase in wheat productivity and iv) search possibilities of simultaneous improvement in grain yield and grain.

## Material and methods

### Source of data

The All India Coordinated Research Project on Wheat and Barley (AICRPW&B) conducts yield evaluation trials to identify wheat genotypes suitable for a particular production environment. The trials are conducted in two trial series i.e. timely-sown (TS) and late-sown (LS) in five wheat zones of the country namely northern hills zone (NHZ), north-western plains zone (NWPZ), north-eastern plains zone (NEPZ), central zone (CZ) and peninsular zone (PZ). This study analysed the data generated by this national wheat research programme of India for the period 2000 to 2020.

arch ppexamined performance of the checks (released varieties) and the new test entries that reached final year of testing (pre-released varieties) during the 20 year period 2000-19.

### Study material and production environments

Study material involved released varieties (checks) and the pre-released high yielding wheat varieties (entries in final year of testing) evaluated in advance varietal trials of AICRPW&B in ten production environments i.e. two production conditions (TS and LS) and five zones. NHZ that covers hills and foothills of the Himalayas has long winter with low temperature while NWPZ and NEPZ represented the Indo-Gangetic plains (IGP). Study material involved released and pre-released high yielding wheat varieties evaluated in two trial series of advance varietal trials constituted by AICRPW&B in five diverse zones of the country i.e. NHZ, NWPZ, NEPZ, CZ and PZ. NHZ that covers hills and foothills of the Himalayas has long winter with low temperature while NWPZ and NEPZ represent the IGP. Among the five zones, NWPZ is the most productive wheat belt of India. Climatic conditions in this zone are most ideal for wheat growth. In comparison, winter is short and climate is normally humid in NEPZ. Wheat crop in CZ often face soil moisture stress and high temperature as climate is hot and dry in this part of India. Peninsula in down south i.e. PZ has similar temperature and soil but climate is not that dry. Planting of timely-sown wheat (TSW) started with the onset of winter and was mostly completed by the end of October in the hills and by the middle of November in the plains. The late-sown wheat (LSW) was planted 15-20 days after the sowing schedule of TSW. Since LSW gets shorter life span therefore short duration genotypes fit in this category. Fertilizer dose in TSW was 150N:60P:40K kgha^-1^ in NWPZ/ NEPZ and 120N:60P:40K kgha^-1^ in the CZ/ PZ and NHZ whereas dosage in LSW was 90N:60P:40K kgha^-1^ throughout the country. No chemical was sprayed while raising the crop under these production environments.

### Variables and statistical analysis

Since the trials involved multiple test sites, the zonal mean of each test entry was computed for plant height (HT), days to maturity (DM), days to heading (DH), 1000 grain weight (TGW) and grain yield. DH denoted the vegetative duration whereas difference between DM and DH represented the grain filling duration (GFD) or the reproductive phase. Standardised data of each environment was computed for regression analysis to assess relationship of NGP’s in grain yield and grain weight. Coefficient of determination (R^2^) derived separately for individual or composite factors highlighted comparative contribution of the NGP’s individually and in combination. Pearson correlation coefficient was calculated to understand association amongst the NGP’s whereas coefficient of variation (CV) was derived to estimate the level of diversity in different parameters. Difference occurring in mean of two populations was compared by “t-test”.

## Results and discussion

### Diversity in production environments

The ten production environments analysed in this study were quite diverse in expression of yield and yield determining attributes (Table 1). Overall performance and the information provided on field expressions during the 21 years revealed significant yield difference between two zones of IGP i.e. NWPZ and NEPZ even when there was no difference in plant height. Difference in height was conspicuous between NHZ and PZ under timely-sown condition as the NHZ crop was 15 cm taller than PZ. NHZ also had 62 days maturity duration advantage in comparison to PZ but there was hardly any difference in wheat productivity and TGW. Overall productivity was highest in NWPZ and CZ and the yield levels also matched in both category of wheat i.e. TSW and LSW even though large maturity difference existed in both categories of wheat. Comparison of reproductive phase very clearly spelt that whatsoever be the difference in maturity duration; time taken to complete grain filling did not differ much. Under timely-sown condition, it was only CZ where grain maturity was completed in 50 days otherwise this process had taken 43-45 days in all other zones. Grain weight in TSW of CZ was exceptionally due to longer reproductive phase. In late-sown wheat, reduction in comparison to timely-sown material was obvious in all variables but zone-wise differentiation was same as observed in TSW. Mean yield of LSW was very low in NHZ mainly because HT was reduced drastically.

**Table 1.** Overall mean and coefficient of variation for six metric traits under diverse production environments of India.

| Characteristic | Timely-sown wheat |  |  |  |  | Late-sown wheat |  |  |  |  |
| --- | --- | --- | --- | --- | --- | --- | --- | --- | --- | --- |
|  | NWPZ | NEPZ | CZ | PZ | NHZ | NWPZ | NEPZ | CZ | PZ | NHZ |
|  | (123) | (136) | (58) | (70) | (95) | (121) | (86) | (89) | (79) | (62) |
| Grain yield (g · m<sup>2</sup>) | 509 | 422 | 492 | 436 | 427 | 416 | 356 | 426 | 386 | 284 |
| Heading days | 96.8 | 79.3 | 67.7 | 61.2 | 126 | 82.6 | 67.7 | 62.2 | 56.4 | 105 |
| Maturity days | 142 | 122 | 117 | 106 | 169 | 121 | 107 | 106 | 97.2 | 145 |
| Grain filling days | 45.6 | 43.0 | 49.6 | 44.4 | 43.5 | 38.3 | 39.6 | 43.7 | 40.8 | 40.0 |
| Plant height (cm) | 93.8 | 91.6 | 88.7 | 82.4 | 98.7 | 88.2 | 85.7 | 80.2 | 79.3 | 82.5 |
| 1000-grain weight (g) | 38.7 | 40.4 | 44.1 | 41.4 | 40.1 | 36.6 | 37.4 | 40.5 | 39.5 | 39.4 |
| *Coefficient of variation (%)* |  |  |  |  |  |  |  |  |  |  |
| Grain yield | 10.2 | 08.9 | 08.4 | 09.8 | 15.1 | 09.8 | 09.0 | 09.9 | 09.1 | 19.8 |
| Heading days | 05.1 | 05.4 | 07.6 | 06.1 | 06.4 | 04.1 | 04.9 | 05.9 | 06.1 | 07.3 |
| Maturity days | 02.8 | 02.7 | 03.3 | 04.2 | 04.5 | 02.4 | 03.1 | 03.1 | 04.1 | 04.6 |
| Grain filling days | 07.3 | 06.5 | 06.5 | 06.3 | 15.8 | 07.5 | 07.5 | 06.1 | 08.8 | 15.3 |
| Plant height | 06.6 | 07.8 | 04.1 | 06.8 | 07.1 | 05.2 | 07.2 | 06.9 | 06.0 | 09.4 |
| 1000-grain weight | 06.4 | 06.4 | 09.7 | 08.2 | 11.2 | 07.6 | 05.9 | 06.1 | 07.2 | 14.1 |

It was evident that wheat productivity did not commensurate with HT and DM in the same manner under varying agro-climatic conditions. Unlike the plains, yield advantage due to HT or DM was hard to realize in the hill region as cold stress might have restricted advantage of HT or DM in TSW. Situation further aggravated in NHZ when there was ceiling on crop duration due to late planting. This kind of abiotic stress led to reduction in HT; consequently, productivity in NHZ-LS environment was lowest in the country. NWPZ and CZ were distinct from other zones in both categories of wheat even though disparity persisted in DM and HT. Even with longer crop duration; GFD in NWPZ was always shorter in comparison to CZ. As a result, improved grain weight compensated the yield loss in CZ which the reduced grain number might have incurred because of shorter vegetative period. Comparison of NWPZ and NEPZ revealed significant differences in maturity and productivity even though there was no big difference in HT. It underlined that association of the NGP’s with wheat yield is based upon the growth conditions prevailing in that particular environment. Influence of NGP’s on wheat productivity cannot be adjudged from differential wheat expression noticed under varying environments. It just helps to understand characteristic features of wheat expression of different production environments.

### Divergence in NGP’s interrelationship

Understanding of relationship amongst the NGP’s is crucial before analysing their impact on wheat productivity. It was observed that besides overall expression, magnitude of variability also differed in study material of each environment (Table 1). There were certain commonalities also in the phenological expressions like CV derived for DM was always less in comparison to DH and GFD in each environment and GFD expressed more variations in comparison to DH and DM in majority of the cases. Maturity differences were lowest in the region where climate was most conducive for wheat growth i.e. NWPZ. NHZ was distinct from the plains as variations in GFD, grain yield and TGW were quite high. DM was less variable in NWPZ under both production conditions. HT variations were highest in LSW of NHZ and lowest in TSW of CZ. It was an indication that relationship between the NGP’s and their association with grain yield could differ in divergent production environments.

Relationship between the NGP’s varied according to the variability noticed in the region. Correlation study revealed strong positive relationship between HT and DM in most of the cases except TSW of NHZ and LSW of NWPZ (Table 2). Although HT and DH are assumed to have strong positive relationship but this association was totally missing in NHZ. This relationship was also not visible LSW of NWPZ. It was obvious that variations in the pre-anthesis period had no significant impact on HT in such environments. DH was correlated positively with DM and negatively with GFD under all conditions but strong association between DM and GFD was not realised in TSW of NWPZ, NEPZ and CZ. It was an indication that variations occurring in DM might not have induced any shift in GFD in such environments. It was clearly evident that if certain associations which are so obvious otherwise (like relationship between HT and DM, DH and HT) fail to establish under certain environments in spite of comparable variation level (CV); it is fair enough to assume that the trend did not exist under those conditions. Every production environment has certain unique NGP relationships which account for differential impact on grain formation and grain development.

**Table 2.** Correlation coefficient between NGP’s under different production environments.

| Relationship | Timely-sown wheat |  |  |  |  | Late-sown wheat |  |  |  |  |
| --- | --- | --- | --- | --- | --- | --- | --- | --- | --- | --- |
|  | NWPZ | NEPZ | CZ | PZ | NHZ | NWPZ | NEPZ | CZ | PZ | NHZ |
| DM~HT | 0.61** | 0.30** | 0.47** | 0.34** | 0.19 | 0.07 | 0.28** | 0.45** | 0.29* | 0.37** |
| DM~DH | 0.74** | 0.76** | 0.78** | 0.78** | 0.62** | 0.59** | 0.59** | 0.71** | 0.55** | 0.63** |
| DM~GFD | 0.09 | 0.01 | -0.05 | 0.56** | 0.39** | 0.33* | 0.45** | 0.27* | 0.59** | 0.29* |
| DH~HT | 0.56** | 0.20* | 0.58** | 0.25* | 0.12 | 0.10 | 0.27* | 0.61** | 0.25* | 0.18 |
| DH~GFD | -0.60** | -0.64** | -0.66** | -0.08 | -0.48** | -0.57** | -0.46** | -0.49** | -0.35** | -0.57** |

\* and \*\* denote significance at $P \leq 0.05$ and $0.01$ , respectively

### NGP relationship with grain yield

Phenology and plant height are major field expressions linked with wheat productivity in any environment, representing manifestation of the genetic (vernalization, dwarfing and photoperiod insensitive genes) and the non-genetic parameters like crop management and weather conditions (Saiyed *et al*., 2009; Yadav *et al*., 2014). Similarly, maturity duration is slated to have strong positive effect on the wheat yield (Garatuza-Payan *et al*., 2018; Singh *et al*., 2017; Xie *et al*., 2015). Another NGP associated with productivity variations is the plant height (Bognár et al., 2007). Since HT is also closely associated with DM, any alteration in height gets reflected in DM and the yield harvest. It is quite obvious that wheat productivity differences under different environments occur mainly because the duration to complete the life cycle differs. Due to diverse production environments and genetic makeup of the test entries; difference in yield and maturity duration are quite large in the Indian wheat. 21-year is a big time frame and so many variations must have occurred climatically and different plant types must have been tested during this long spell in every production environment. Fluctuations in weather conditions and diversity in the tested material therefore must have recorded different levels of variations in the grain as well as non-grain attributes (Table 1).

It was amply clear that NGP’s also play important role in regulating the yield potential as each one expressed significant relationship in 7-8 out of 10 environments (Table 3). In view of differential relationship amongst the NGPs, their contribution also varied under diverse production environments. Regression analysis revealed that magnitude of association between NGP’s and grain yield varied in each environment. It was amply clear that NGP’s also play important role in regulating the yield potential as each one expressed significant relationship in 7-8 out of 10 environments. Amongst all environments, it was only TSW of NWPZ where every NGP established significant relationship with grain yield. It simply means that when growth conditions are favourable in a given environment, number of NGP’s associated with yield also popup. There was no NGP which could establish relationship with yield under all conditions and the least important amongst them was GFD.

**Table 3.** Relationship of individual NGP with wheat yield in different production environments of India.

| Parameter | Coefficient of determination ($R^2$) | | | | | | | | | |
| --- | --- | --- | --- | --- | --- | --- | --- | --- | --- | --- |
|  | Timely-sown wheat |  |  |  |  | Late- sown wheat |  |  |  |  |
|  | NWPZ | NEPZ | CZ | PZ | NHZ | NWPZ | NEPZ | CZ | PZ | NHZ |
| HT | 0.45*** | 0.09*** | 0.14** | 0.03 | 0.00 | 0.08** | 0.04* | 0.12*** | 0.10** | 0.20*** |
| DH | 0.09** | 0.01 | 0.10* | 0.15*** | 0.04 | 0.03 | 0.10** | 0.10** | 0.16*** | 0.08* |
| GFD | 0.09** | 0.06** | -0.09* | 0.00 | 0.01 | 0.07** | 0.02 | 0.03 | 0.00 | 0.03 |
| DM | 0.39*** | 0.11*** | 0.03 | 0.12** | 0.07** | 0.22*** | 0.18*** | 0.23*** | 0.09** | 0.03 |
| HT+DH | 0.46*** | 0.09** | 0.16** | 0.16** | 0.04 | 0.11*** | 0.12** | 0.13** | 0.21*** | 0.25*** |
| DH +GFD | 0.45*** | 0.18*** | 0.12* | 0.16** | 0.07** | 0.23*** | 0.19*** | 0.24*** | 0.16*** | 0.08 |
| All NGP's | 0.63*** | 0.22** | 0.17** | 0.16** | 0.08* | 0.30*** | 0.20*** | 0.27*** | 0.21*** | 0.27*** |

\*, \*\* and \*\*\* denote significance of $R^2$ at $P \leq 0.05$ , $0.01$ and $0.001$ , respectively

Although wheat productivity in NWPZ matched CZ under timely-sown condition, huge difference could be seen in the impact of NGP’s. It underlined that the variations created in NGP’s through scientific interventions and natural climatic variations were exploited to high capacity in TSW of NWPZ whereas prospects of exploiting such variations were rather limited in CZ (Table 1 and 3). Relevance of individual NGP in yield of TSW in NWPZ was very high in case of HT and DM (R^2^: ≥ 0.40). Highest R^2^ value recorded in TSW of all other zones was 0.15 noticed for DH in PZ. In LSW however, R^2^ ≥ .20 could be noticed for HT in NHZ and DM in NWPZ/ CZ.

Height and heading are two important growth factors related to the vegetative phase and it is important to understand and fragment their contribution during this developmental phase. Composite impact of these two factors was similar to that of HT in both production conditions of NWPZ, TSW of NEPZ/ CZ and LSW of NHZ (Table 3). It ascertained that during vegetative phase, it was only the height (not heading) which regulated wheat yield in such environments. It means that when HT shows sign of increase due to favouring climate or the scientific interventions; no big yield benefit is realised through DH in such environments which means that the grain number remains nearly the same. Significant increase in grain number under such situations is feasible only when tillering is enhanced through breeding. But when lateral growth intensifies, flowering is automatically delayed and the prolonged vegetative phase results in increased grain number. At the same time even if flowering gets extended due to agri-ecology or genotypic differences but crop face abiotic pressure in the vegetative phase, there can always be reduction in plant height which ultimately results in the yield loss (Anjum *et al*., 2017). On the contrary, out of the two only DH expressed significant contribution in TSW of PZ and LSW of NEPZ/ CZ which showed that benefit of congenial climatic conditions in these environments was realised through DH only. Role of NGP’s in grain yield was limited in NHZ as only DM mattered to some extent in TSW whereas only HT was prominent in LSW. Contribution of vegetative phase linked NGP’s was significant in LSW of PZ condition and synergy was derived when HT and DH were regressed together against yield.

Although several reports point relevance of GFD in grain development (Xie *et al*., 2015; Wu *et al*., 2018), its positive contribution in yield could only be cited in some production environments of northern plains of India, especially TSW of NWPZ and NEPZ; and LSW of NWPZ. Synergy was also visible when DH and GFD were regressed together in both zones of IGP as DM turned highly significant even though individual impact of HD or GFD was not high. Results related to CV had illustrated that variation level recorded in DM was magnified in DH and GFD in many environments and this distinction was very clear in northern India (Table 1). It underlines that increase in DH can lead to better grain bearing in northern India but higher yield gain can only be achieved when proper GFD is available. Similarly, enhanced GFD might fail to deliver good yield if flowering is enforced early. Similar relationship could also be noticed in LSW of NEPZ and CZ. CZ was a unique example where yield in TSW was benefitted by DH but impact of GFD was negative. As result, yield registered no significant relationship with maturity in this particular environment. Increased TGW in TSW of CZ was at the cost of reduced grain number affected by shortening of the vegetative phase. This was the only production environment in the country where TGW was adversely related to wheat productivity. Further, irrelevance of a given NGP in grain yield cannot be attributed to lack of variability (Table 1); it can also happen when direct effect of a given component is marginal.

### Collective contribution of NGP’s in grain yield

In multiple regression analysis, R^2^ value obtained through combinations like HT+DH+GFD, HT+DH+MAT and all 4 NGP’s together was similar. It’s only because GFD was a derived component from DM and DH. Since, it’s not easy to exercise selection on the basis of GFD in the field, this factor was excluded and the choice was limited to HT, DH and DM. Collective impact of NGP’s was highly significant in wheat yield of each production environment (Table 4). Indirect contribution of NGP’s was significant in every environment but the magnitude (R^2^ value) varied from 0.07 to 0.61. In comparison to individual effect, combination of NGP’s was beneficial in majority of the cases. This impact was highest in NWPZ in both categories of wheat and lowest in TSW of NHZ. R^2^ value underlines percent variations in yield associated with a given NGP or group of NGP’s. In most congenial wheat growing environment of the country i.e. TSW of NWPZ; even 61% yield variations could be accounted by NGP’s alone. In contrast, their contribution was limited to just 7%. Actually level of cold stress vary each year in the hill due to climatic fluctuations which results into high degree of yield fluctuations (Table 1). In all other production environments, 16 to 30% variations in grain yield were accrued through NGP’s. High R^2^ value does not indicate that TSW of NWPZ could make best use of the climatic conditions for suited for wheat growth. It could also have happened because of the desired genetic variations created in NGP’s through wheat breeding. Variations derived through the scientific interventions can be spotted in the wheat genotypes developed in this region for height and the phenological expressions. Emphasis in this region is also given to effective tillering which influence height, heading and maturity as well. Plotting of grain yield against maturity period countrywide (N: 919) had clearly illustrated that NWPZ-TS environment is distinct, indeed as entries developed and tested in this environment (maturity: 142 ± 4 days) formed a separate cluster altogether (Fig 1). Entries of the maturity range noticeable in NHZ (Table 1) were found scattered in another cluster. Irrespective of the production condition, all test entries pertaining to the Indian plains (except NWPZ-TS), were accommodated in single cluster.

**Table 4.**
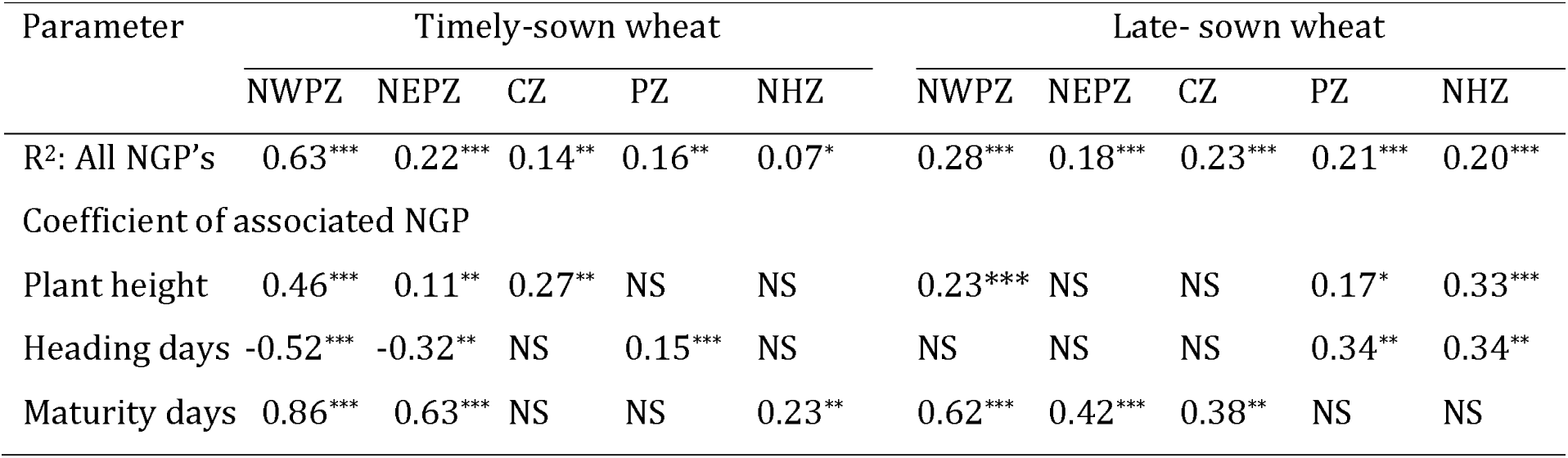
Multiple regression statistics of key contributors in wheat yield in different production environment.

**Figure 1.**
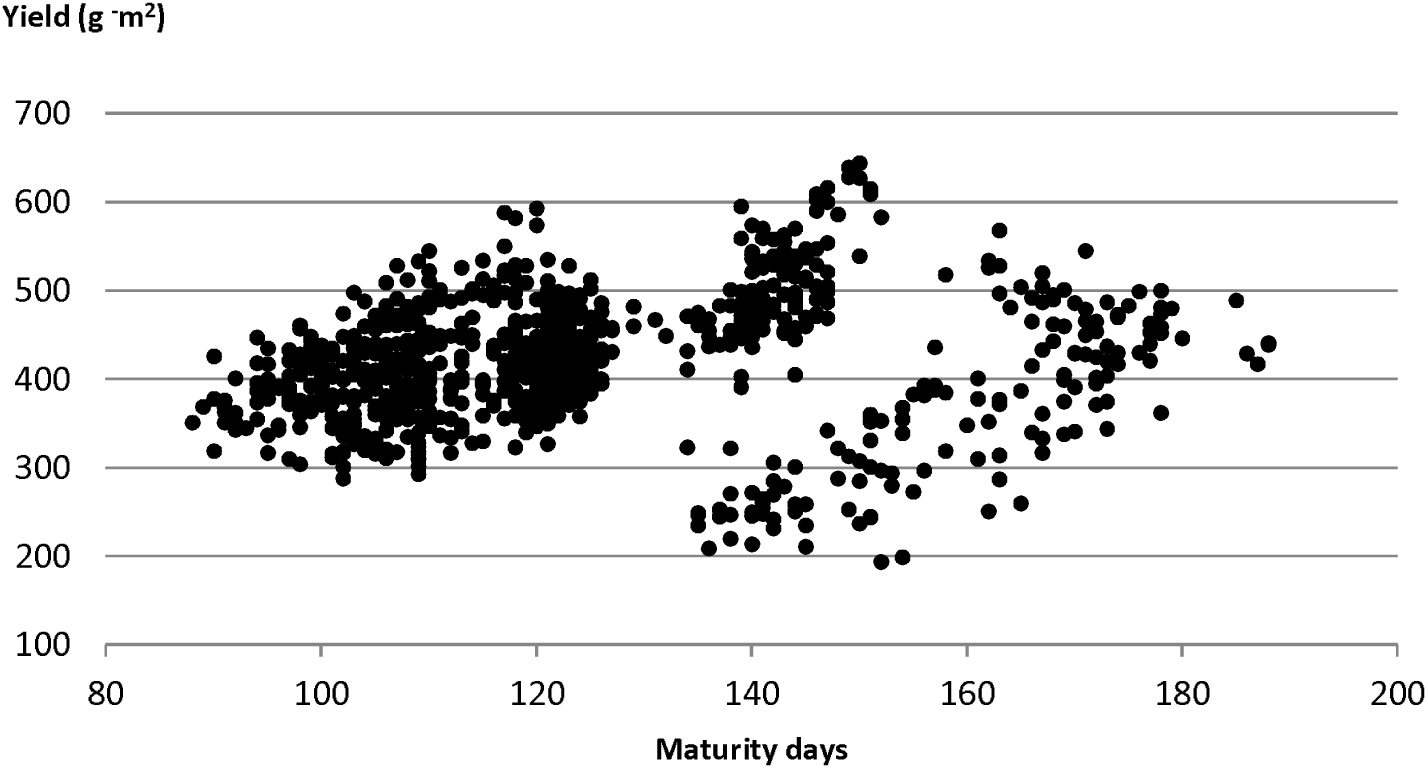
Maturity duration and wheat productivity under Indian environments.

### Key contributors in wheat yield

Impact of individual components was also examined in the multiple regression analysis. It was observed that all of them expressed significance only in TSW of IGP. Otherwise just one or two were main derivers of this relationship. Analysis was further done to keep only those constituents which exerted significance in multiple regression analysis (Table 4). Even though individual impact of each constituent (HT, DH and DM) was positive in IGP (Table 3), contribution of DH turned negative in this combined relationship. In NEPZ-TS environment, only HT and DM were relevant individually (Table 3) but regression equation indicated that just like the adjoining zone i.e. NWPZ, DH was also equally important and its indirect contribution was significant, too. Under late-sown condition, just two factors i.e. HT and DM were found relevant in NWPZ whereas just one figured in NEPZ. In CZ, only height mattered in TSW whereas maturity was crucial for LSW. Situation was different in the adjoining PZ as only DH was the real deriving force in TSW whereas HT and DH were important in LSW. In NHZ, DM proved to be the main yield regulator of TSW whereas HT was key yield determinant in LSW. Importance of height in LSW of NHZ was also evident when zone-wise comparison was made earlier in Table 1.

### Selection strategy to harness yield and grain weight through NGP’s

Genetic variability, climatic variations and direct effect; they all matter in deciding the bottleneck factor in grain yield under any production environment. It was quite evident in this study that NGP’s can be exploited as yield contributor in wheat. For effective implementation, it is imperative to devise a strategy based upon minimum number of NGP’s in such a way that there is simultaneous gain in yield and TGW. If not, at least there should not be a case where yield gain is harnessed with a premium on grain weight. Since all NGP’s are not related to yield in every production environment (Table 3), only the key components can be exploited to formulate selection index. Enrichment of wheat yield through enhanced plant height and prolonged vegetative duration had been suggested for the Indian subcontinent (Jamali and Ali, 2008; Laxman *et al*., 2014). Reports from Pakistan and China had also emphasised selection through improved height and larger flowering or reproductive periods (Duan *et al*., 2018; Khan *et al*., 2000; Wu *et al*., 2018). Advantage of height and crop phenology had also been reported in some Indian environments by Mohan *et al*. (2017). Such phenotypic expressions are indicators of good biomass accumulation accrued from healthier vegetative growth. In the era of green revolution, reduced height had been preferred in wheat breeding for a long time due to less lodging loss but it put a ceiling on the plant height, therefore, the role of tall-dwarfs had also been acknowledged in the development of the high-yielding semi-dwarf wheat’s in the green revolution (Würschum *et al*., 2017; Mohan *et al*., 2017; Mohan *et al*., 2017). It is also well established that flowering in wheat depends upon aaccumulation of certain amount of heat units and this had been amply demonstrated earlier in two contrasting zones of India i.e. NWPZ and CZ (Mohan *et al*., 2017a). Thus breaking the yield barrier in wheat will require fine tuning of the sink-source relationship which requires simultaneous increase in grain number and the available assimilates for grain filling. But making selection for high tillering or grain bearing is not easy. Often such plants flower late and get less time for proper grain filling.

On the basis of information gathered about the key components (Table 4), simple and easy to adopt inference can be generated for simultaneous improvement in grain yield and grain weight. Identification of key NGP’s make the job easy for the breeders as by application of these 1-2 indicators in the field, significant yield improvement can be anticipated in wheat. The key factors were common in TSW of NWPZ and NEPZ which underlines that positive selection for height and maturity in early flowering genotypes can be highly useful to improve productivity of timely-sown wheat in the IGP (Table 5). Besides yield, this selection strategy can also benefit TGW in a significant way. To develop high yielding genotypes for LSW of NWPZ; field selection based upon just two NGP’s i.e. HT and DM can be instrumental for significant productivity improvement but yield gain will be less in comparison to NWPZ and there shall not be any improvement in TGW either. In LSW of NEPZ, only DM is the main predictor for yield and it also helps to improvise TGW as well. Plant height should be given maximum importance in TSW of CZ whereas maturity is crucial for LSW of the region. However, no extra advantage through TGW is expected through this planning. Selection tool can be different in the adjoining PZ where DH is the lone predictor for grain yield in TSW. When coupled with HT, significant yield gain can also be anticipated in LSW of the region. Just like CZ, there is hardly any chance of grain size improvement through NGP’s in PZ as well. Since yield variations are usually high in NHZ, precision might lack in the estimated benefits of NGP’s. Still, selection for prolonged maturity in TSW and plant height in LSW can be expected to boost wheat productivity. Preference to height in LSW of NHZ will also come in aid to grain weight of the harvested produce.

**Table 5.** NGP based selection criteria for simultaneous improvement in yield and grain weight of wheat.

| Zone | Selection criteria | Coefficient of determination (R<sup>2</sup>) |  |
| --- | --- | --- | --- |
|  |  | Grain yield | Grain weight |
| *Timely-sown wheat* |  |  |  |
| NWPZ-TS | Height, early flowering and maturity | 0.631*** | 0.303*** |
| NEPZ-TS | Height, early flowering and maturity | 0.217*** | 0.130*** |
| CZ-TS | Height, early flowering | 0.141** | 0.058 |
| PZ-TS | Days to heading | 0.154*** | 0.022 |
| NHZ-TS | Early flowering, maturity | 0.074* | 0.004 |
| *Late-sown wheat* |  |  |  |
| NWPZ-LS | Height and maturity | 0.283*** | 0.034 |
| NEPZ-LS | Maturity | 0.184*** | 0.178*** |
| CZ-LS | Maturity | 0.234*** | 0.005 |
| PZ-LS | Height and days to heading | 0.206*** | 0.065 |
| NHZ-LS | Height | 0.202*** | 0.121** |

This study further adds that height accompanied with unstressed maturity duration ensures higher wheat productivity in most favourable wheat growth environment in India i.e. NWPZ-TS. Under late-sown condition, it is often difficult to pick genotypes which mature late as elevated temperature conditions and the hot winds enforce senescence in the leaves. Nevertheless, it is also well known that Sonalika, a prominent old cultivar for late-sown condition, had expressed productivity level well below the present varieties mainly because the new high-yield varieties have comparatively longer maturity duration. During last stage of this study period, all these components had been exploited in variety development programme of NWPZ-TS. Annual progress report of crop improvement (ICAR-IIWBR, 2019) had mentioned tremendous improvement in grain yield, plant height, bio-mass accumulation, harvest index, heading days and maturity duration when growth condition were highly conducive. If performance of four leading TSW varieties of NWPZ (HD 2967, HD 3086, DBW 88 and WH 1105) common during the crop seasons 2018 to 2020 is compared, average yield harvest was highest in 2019 (612 g ^−^m^2^)) in comparison to the previous year when yield was restricted to 539 g ^−^m^2^. It happened because there was 10 days increase in maturity duration (from 141 to 151 days) and 6 cm increase in plant height (from 98 to 104 cm). With increased height and favouring phenology, not only yield but TGW also increased from 38.8 to 40.3 g. In comparison to 2019 harvest season, wheat productivity in this region declined to 580 g ^−^ m^2^ in 2020 as height was reduced by two cm and maturity period by three days. Comparison of NGP’s further revealed that genetic differences also counted for differential expressions in these four high yielding varieties. HD 3086 excelled because of longer grain ripening period (49 days) and good plant height (101 cm). HD 2967 drew advantage of extra height (103 cm) and longer maturity duration (148 days). DBW 88 had plant height (102) and maturity duration (147 days) almost similar to HD 2967 but it had advantage of early flowering and prolonged GFD. In spite of shorter maturity period (145 days) and reduced plant height (99 cm), WH 1105 was high yielding because phenology was partitioned well (HD: 98 days, GFD: 47 days).

## Conclusion

Diverse production environments necessitate special wheat improvement strategy. At a time when wheat is touching the yield plateau, it is pertinent that vista of non-grain plant attributes is also reviewed. Breeders do keep an eye on these aspects while exercising selection in the segregating material but which parameter is to be emphasized in a given environment is the key. Artificial intelligence gathered through this investigation offers some silver lining. Simple and easily adoptable selection methodology devised through this study will surely enhance the prospects of yield improvement further. This information on potential uses of the non-grain yield determinants can bridge some gap in the yield barrier realised not only in India but all over the world. It assures that with some strategic planning; prospects of wheat productivity enhancement can be improvised without directly touching the grain related parameters. Knowledge gained from this 21-years study period assures that variability in NGP’s exists in each production environment. In the past, this variation was exploited to certain extent unknowingly. But when applied with some strategic planning, prospects of yield improvement can surely be improved further. It’s easy to make selection for these phenotypic traits in the field as it will improvise biomass through plant height; grain number through enhanced vegetative phase and grain weight by adjusting the grain filling duration. Since impact can be different under divergent environments, breeders can choose the factors or combination of NGP’s required for yield improvement in a given environment.

## Acknowledgments

The work is an outcome of a core project funded by ICAR (Project No. CRSCIIWBRCIL201500100182), New Delhi and the authors express their sincere thanks to the Director, ICAR-IIWBR for permitting use of the data generated in All India Coordinated Research Project (AICRP) on Wheat and Barley for this analysis. The efforts made by associated wheat research workers in trial conduct and data reporting are also acknowledged.

